# The Effects of Migration and Assortative Mating on Admixture Linkage Disequilibrium

**DOI:** 10.1101/056168

**Authors:** Noah Zaitlen, Scott Huntsman, Donglei Hu, Melissa Spear, Celeste Eng, Sam S. Oh, Marquitta J White, Angel Mak, Adam Davis, Kelly Meade, Emerita Brigino-Buenaventura, Michael A LeNoir, Kirsten Bibbins-Domingo, Esteban G Burchard, Eran Halperin

## Abstract

Statistical models in medical and population genetics typically assume that individuals assort randomly in a population. While this simplifies model complexity, it contradicts an increasing body of evidence of non-random mating in human populations. Specifically, it has been shown that assortative mating is significantly affected by genomic ancestry. In this work we examine the effects of ancestry-assortative mating on the linkage disequilibrium between local ancestry tracks of individuals in an admixed population. To accomplish this, we develop an extension to the Wright-Fisher model that allows for ancestry based assortative mating. We show that ancestry-assortment perturbs the distribution of local ancestry linkage disequilibrium (LAD) and the variance of ancestry in a population as a function of the number of generations since admixture. This assortment effect can induce errors in demographic inference of admixed populations when methods assume random mating. We derive closed form formulae for LAD under an assortative-mating model with and without migration. We observe that LAD depends on the correlation of global ancestry of couples in each generation, the migration rate of each of the ancestral populations, the initial proportions of ancestral populations, and the number of generations since admixture. We also present the first evidence of ancestry-assortment in African Americans and examine LAD in simulated and real admixed population data of African Americans. We find that demographic inference under the assumption of random mating significantly underestimates the number of generations since admixture, and that accounting for assortative mating using the patterns of LAD results in estimates that more closely agrees with the historical narrative.

## 2 Introduction

One of the most common assumptions in human population genetics analyses is that of Hardy-Weinberg Equilibrium (HWE). The HWE assumption in turn enforces a set of additional conditions including the absence of selection, infinite population size, and importantly, random mating. We and others have shown that assortative mating is a common phenomenon[1, 2, 3] and many phenotypes including height, education level, and personality traits are correlated between spouses [4]. For Latinos and other admixed populations the African, Native-American, and European proportions of individual’s genomes can be correlated between spouses. We recently demonstrated that the genomic ancestry of Latino couples is highly correlated [1], and refer to this as ancestry-assortative mating. Thus, the assumption of random mating and therefore Hardy-Weinberg Equilibrium is not satisfied in practice, and the implication of this observation for population and evolutionary genetic studies remains unclear.

The assumption of random mating is used in many types of population and quantitative genetics analyses. Particularly, random mating is assumed both in analysis of population genetics data and when inferring population parameters such as recombination rates, mutation rates, selection, heritability, and others. Moreover, methods for quality control and data cleaning often make the random mating assumption. For example, methods for haplotype phasing typically compute the likelihood of the genotype as the product of the likelihoods of each of the haplotypes, and this derivation is based on the random mating assumption[5]. Similarly, such likelihood derivations are also common in methods for the inference of identity by descent and inference of ancestry from genomic data[6]. Thus far, the sensitivity of these methods to the assumption of assortative mating has not been evaluated. In principle, realistic violations of the random mating assumption may not be detrimental to existing methods, however this needs to be taken to the test.

In this paper we explore the robustness of specific genetic features and their inference from genetic data to assortative mating. Because assortative mating in Latinos has been shown to be affected by ancestral proportions, we focused our analysis on the behavior of ancestry linkage disequilibrium under assortative mating. We propose a random generative model for population dynamics under assortative mating which is due to population structure. Our model follows the spirit of the Wright-Fisher model, and makes the assumption that the correlation of ancestry proportions between spouses stays fixed across generations. Particularly, when the correlation of ancestry proportions is zero, our model is equivalent to the Wright-Fisher model.

We develop mathematical theory that describes the decay of local ancestry disequilibrium (LAD) as a function of assortative mating strength, migration rate, recombination rate, and the number of generations since admixture began. Thus, one can use these results in order to infer the demographic history of admixed populations. Several methods for demographic inference in admixed populations exist including ones that use patterns of LD decay [7], local ancestry track length distribution [8], and the distribution of identity by descent segments[9]. However, these methods assume random mating, and under assortative mating LD decay follows a different pattern [10]. Using simulations we demonstrate that our mathematical derivation matches empirical LAD decay. Furthermore, we develop the theory with migration rates from the ancestral populations, and we demonstrate that in the presence of assortative mating, one may erroneously conclude that there has been active migration, and vice-versa.

We applied our analysis to a dataset of 1730 African Americans from the Study of African Americans, Asthma, Genes and Environments (SAGE) study[11]. We first used ANCESTOR [12] to show that the correlation of African ancestry between the spouses in the last generation is approximately 0.32. We then used our analysis to infer the number of generations and migration patterns in the African American population. Under the assumption of no migrations and random mating, an analysis of LAD resulted in an estimated of the number of generations since admixture of 3. Adding assortment and migrations we find that the estimated number of generations since the admixture event is 15. Assuming a generation time of 25 years this places the initial migrations in the mid 17th century which is consistent with the history of African Americans[13].

## 3 Methods

**The model** We assume the following alternative to Wright-Fisher. Let *N* be the number of individuals in each population (effective population size divided by 2). Each individual has two haplotypes, so the total number of haplotypes is 2*N*. Also, we assume the population is a recently admixed population with two ancestral populations (referred to as population 1 and population 2), and let *θ*_*i*_ denote the fraction of the genome with population 1 ancestry.

In the next generation, each individual picks two parents from the current generation, such that the correlation between the ancestry of the two parents is a fixed value *P*. One way of generating such mating in silica is the following. We randomly pick the set of mothers (with or without replacements) from the original distribution. We then randomly choose the set of fathers (with or without replacements). Now, for each of the parents we give a score *score*_*i*_ = *θ*_*i*_ + *ϵ*_*i*_, where *θ*_*i*_ is the global ancestry of the parent, and *ϵ*_*i*_ is drawn from a normal distribution *N*(0,σ^2^). We then sort the mothers and the fathers based on their score and we let the mother with *i* − *th* largest score marry the father with the *i* − *th* largest score. We then compute the correlation between *corr*(*θ*_*m*_, *θ*_*f*_), where *θ*_*f*_, *θ*_*m*_ are the ancestries of the mother and the father. We choose *σ* such that *P* = *corr*(*θ*_*m*_, *θ*_*f*_). We note that our analysis below does not rely on this specific procedure; particularly, the distribution of parents for the new generation can be quite general, and our only assumption is that *P* is constant across the generations. Note that this assumption may seem restrictive at first, however the case of random mating is far more restrictive, since there one requires that *P* = 0 in all generations.

**Local Ancestry disequilibrium** Denote by 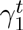 the probability of having an allele from ancestry 1 at a given position at generation *t*. Furthermore, for a pair of positions, let 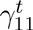 denote the probability of having an allele from ancestry 1 at the two positions. We define a new statistic, termed local-ancestry linkage disequilibrium, denoted by *LAD*. We define 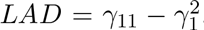. We are interested in the expected value of *LAD*^*t*^ (*LAD* at generation *t*) as a function of the recombination rate *r*, the number of generations *t*, and the original local ancestry linkage disequilibrium *LAD*^0^.

For the following derivation, we will assume that the population size is infinite. We will later show that empirically this assumption does not have a substantial effect for realistic values of *N*. We will first assume that there is no migration and we will relax this assumption in the next section.

Since there is no migration and the population size is infinite, the mean of *θ* is fixed across the generations (remember that the marginal distribution of the mothers and the fathers is the same and is simply a random draw from the current generation)[14]. We denote *μ* = *E*[*θ*]. Let *V*_*t*_ = *Var*(*θ*_*t*_) be the variance of *θ* in generation *t*, and let *ρ*_*t*_ = *PV*_*t*_ be the covariance *ρ*_*t*_ = *cov*(*θ*_*m*_, *θ*_*f*_). For *t* > 1 we have:

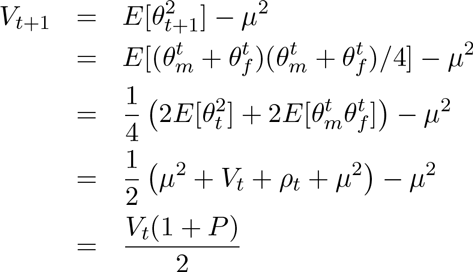

This demonstrates that the variance of genome-wide ancestry is larger when there is assortative mating. Now, we know

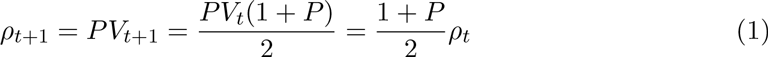

Note that for *t* = 0, *ρ*_0_ = *V*_0_ since there was no assortative mating prior to the admixture event, and therefore for *t* = 1 the above calculation gives *V*_1_ = *V*_0_, and *ρ*_1_ = *PV*_0_ = *P*_ρ0_. In order to simplify the notation, we change the indices, so that generation *t* = −1 corresponds to the time of encounter of the two population and *t* = 0 is the first generation after admixture. Therefore, we have that Equation 1 holds for every *t* ≥ 1.

We now find a recursion formula for *LAD*^*t*^. Let *r* be the probability for an odd number of recombinations between the two positions in a given meiosis. Hence,

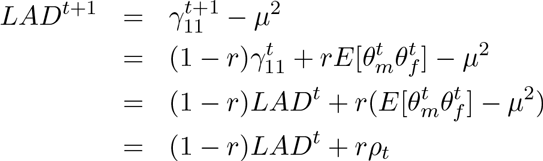

We are now ready to describe our main result:

#### Lemma 3.1

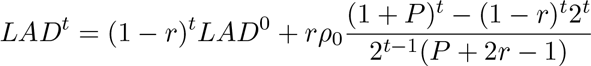

**Proof** We show this is true by induction. It is easy to verify that since *LAD*^1^ = (1 − *r*)*LAD*^0^ + *r*_*ρ*0_, the base case *t* = 1 holds. Assume the lemma holds for *t* and we will prove it for *t* + 1.

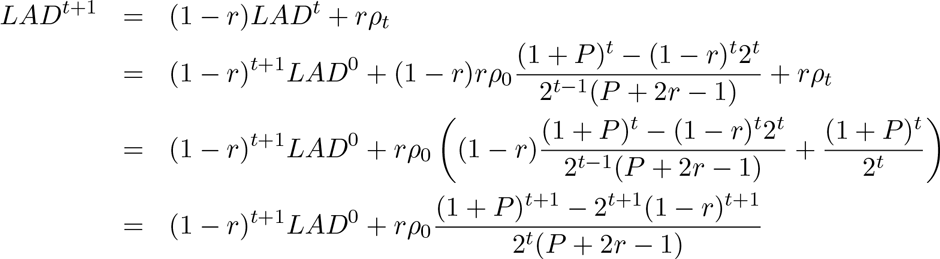

**LAD under migration.** We now assume that in each generation a fraction *m*_1_ of the population is replaced by individuals from the first population (*θ* = 1), and a fraction *m*_0_ of the population is replaced by individuals from the population *θ* = 0. We denote by *m* = *m*_1_ + *m*_0_, and 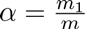. Since there is migration, the mean global ancestry is changing over time, and we let *μ*_*t*_ = *E*[*θ*_*t*_] the average values of *θ* when an individual is randomly sampled from the population. For simplicity of notation, we denote *x*_*t*_ = *μ*_*t*_ − *α*, and we note that *x*_*t*_ is exponentially decreasing. Since *μ*_*t*+1_ = *αm* + (1 − *m*)*μ*_*t*_, we have that *x*_*t*+1_ = (1 − *m*)*x*_*t*_ and therefore *x*_*t*_ = *x*_0_(1 − *m*)^*t*^.

We now show the following lemma:

#### Lemma 3.2

*If there is a sequence y_0_,y_1_,…, satisfying the recursion equation y_t+1_* = 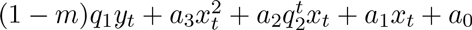, *then*

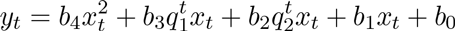

**Proof** To prove the base of the induction we need to satisfy 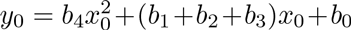, which is a simple linear equation. We will show that the induction step adds two more linear equations. Assume the lemma holds for *t*, and consider *y*_*t*+1_:

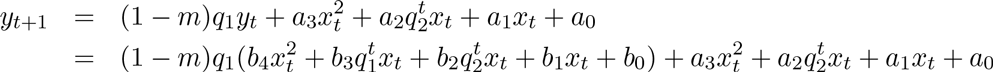

Now, note that *x*_*t*+1_ = (1 − *m*)*x*_*t*_. Therefore:

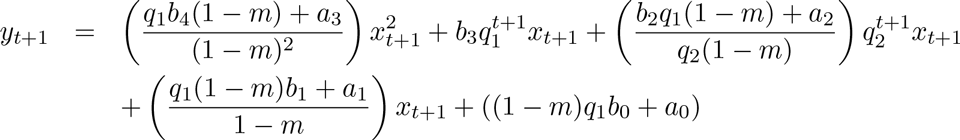

We now set:

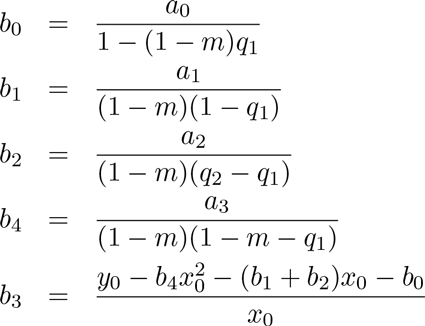

Next, we observe:

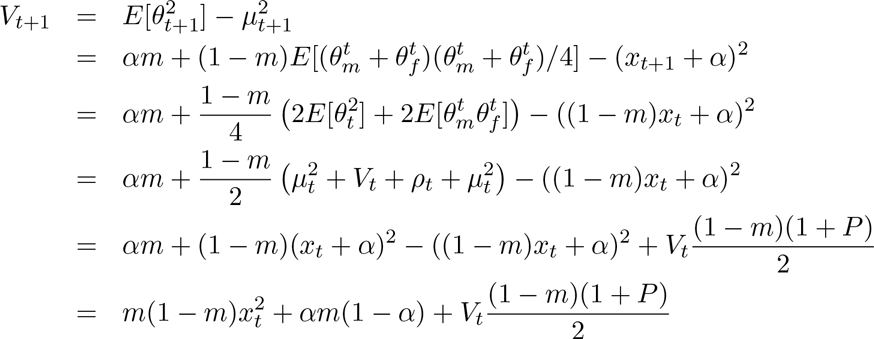

By Lemma 3.2, we have 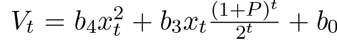, for *b*_4_, *b*_3_, *b*_0_ specified in the lemma. Note that based on the lemma’s proof, *b*_1_ = *b*_2_ = 0. Now,

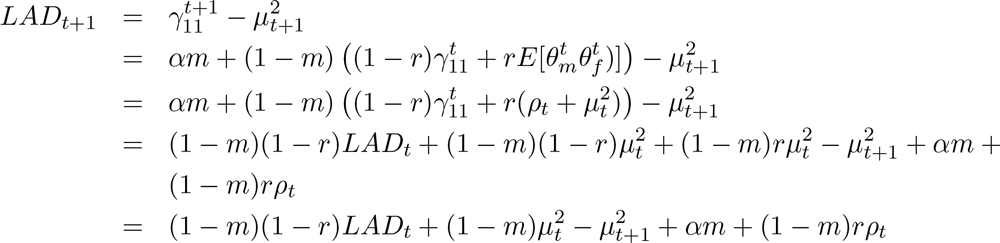

Therefore, noting that *μ*_*t*+1_ = (1 − *m*)*x*_*t*_ + *α*, we have

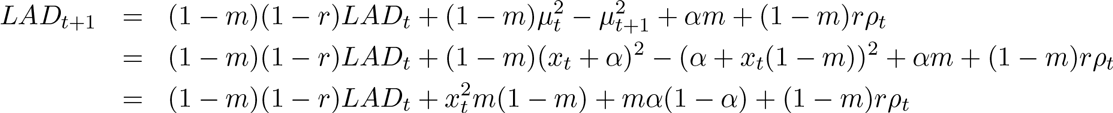

Now, recall 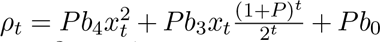. Therefore, we have the form 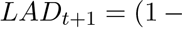 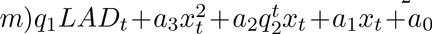 satisfying Lemma 3.2 with the following values:

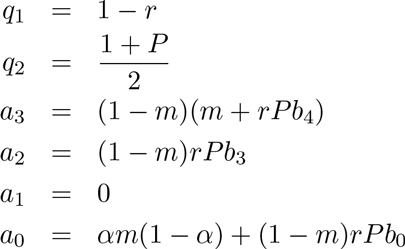

Thus, for *c*_0_, *c*_*1*_, *c*_2_, *c*_3_, *c*_4_ taken from Lemma 3.2 we have

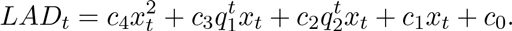

Plugging in the values of *q*_1_, *q*_2_, and the fact that *x*_*t*_ = *x*_0_(1 − *m*)^*t*^, we get

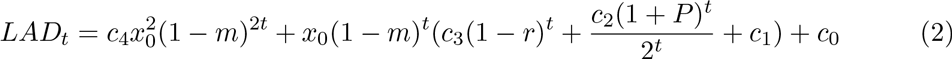

## 4 Results

When applied to the genome, we can estimate the value of LAD for known values of *r* by averaging the observed LAD across the genome. We can now fit the values of *m*, *t*, and *P* based on the distribution of the LAD as a function of *r* in the current generation. It is therefore important to understand the dependency of the distribution of LAD for varying values of *r* as a function of *t*, *P*, and *m*. In what follows, we explore the behavior of LAD under different settings.

We first consider the case where *m*_1_ = *m*_2_ = 0, i.e., there is no migration, and *P* = 0.6. In Figure 1 we observe that there is a clear separation between the different curves for the different numbers of generations since admixture, and it should therefore be easy to estimate the time of admixture event under the assumption of no migration and *P* = 0.6.

**Figure 1:**
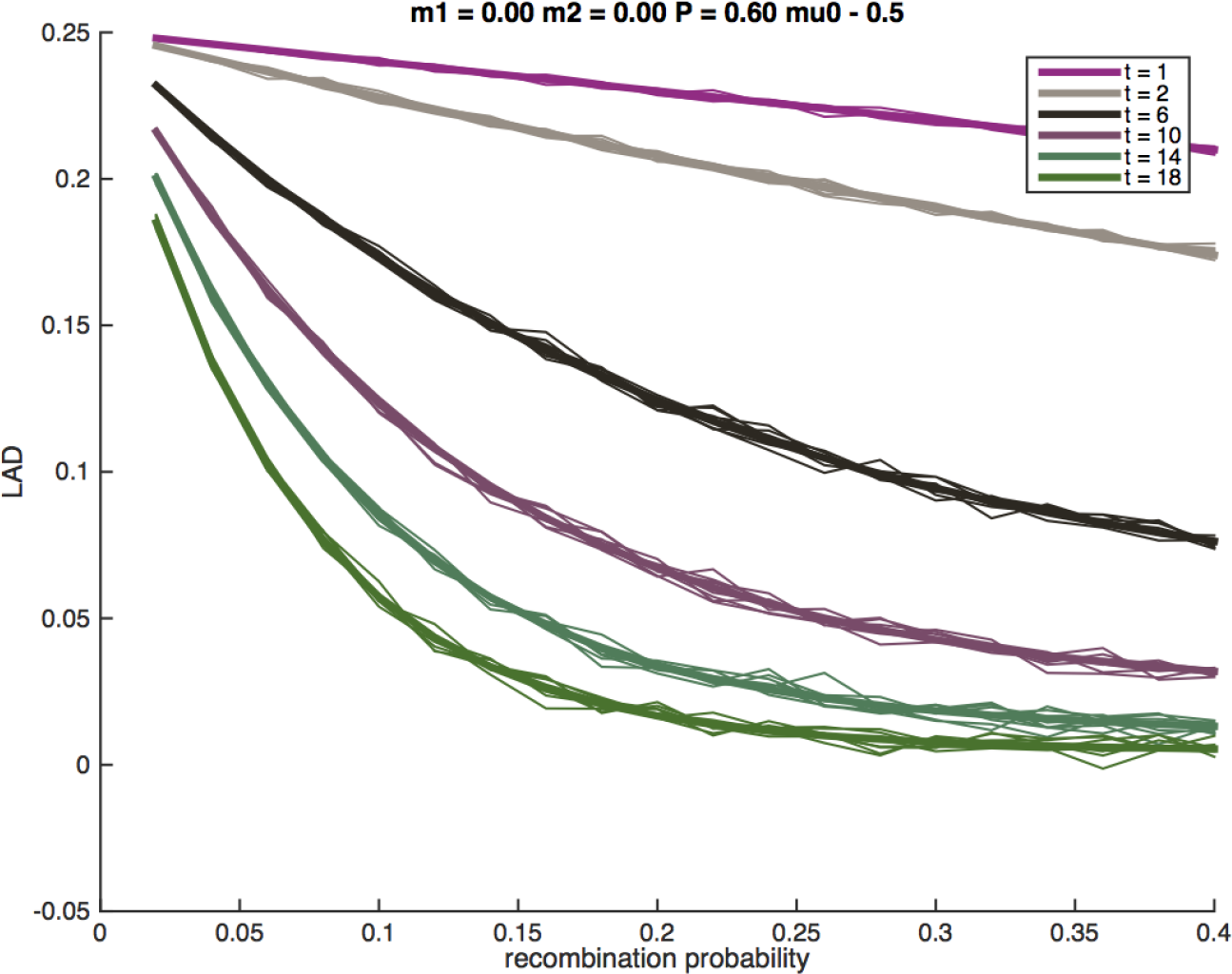
The distribution of LAD for different values of t with no migration(and *P* = 0.6). The thick lines correspond to the expected LAD based on Lemma 3.1, and the thin lines correspond to simulation runs of a single locus in the genome.

Next, we study the effect of *P* on the LAD distribution. In Figure 2 we plot the LAD distribution under no migration, after 10 generations of admixture, with varying values of *P*. Evidently, strong assortative mating with large values of *P* results in a substantially different levels of LAD. However, we observe that low values of *P* are harder to distinguish, and therefore we expect that random mating is a robust assumption for any statistic that uses LAD or its derivatives, as long as assortative mating is weak (e.g., *P* < 0.5).

**Figure 2:**
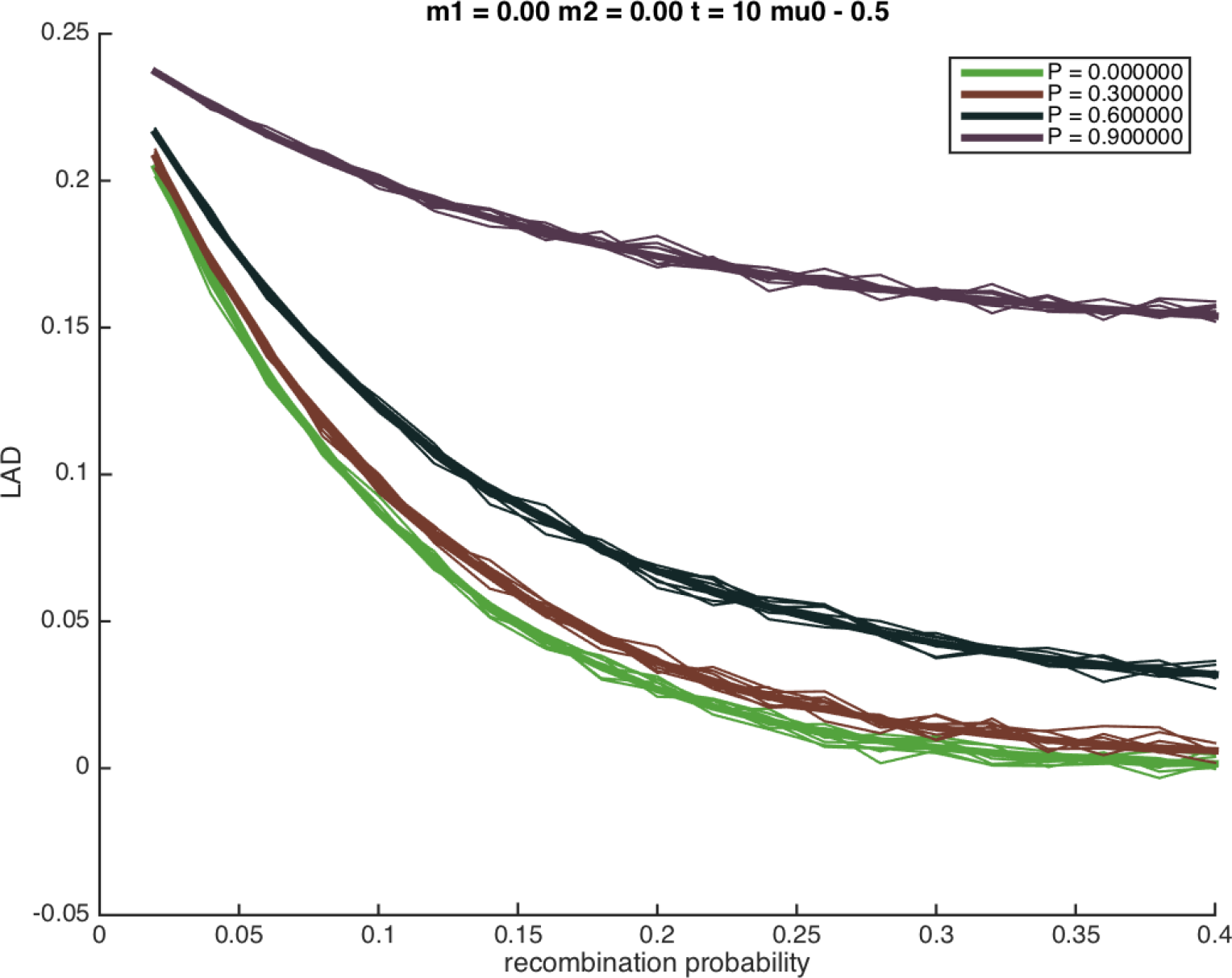
The distribution of LAD for different values of *P* with no migration. The thick lines correspond to the expected LAD based on Lemma 3.1, and the thin lines correspond to simulation runs of a single locus in the genome.

Since typical analysis of genetic data assumes random mating, we attempted at understanding the potential risk in making the assumption in the presence of assortative mating. Thus, we consider the case where there is assortative mating, and we try to estimate the time of admixture under the assumption of random mating. For ancient admixture the difference between the estimates under assortative mating and random mating is not substantial (about 10% - data not shown). For recent admixture (10-20 generations), we observe that there is a considerable difference between the true LAD curve compared to the LAD curve under random mating, and moreover, the true LAD curve is similar to LAD curves that assume random mating but that are substantially more recent. Specifically, in Figure 3 the admixture event occurred 10 generations ago under a strong assortative mating (*P* = 0.8), however under random mating, the LAD curve that corresponds to *t* = 4 is the most similar to the true LAD curve. In Figure 4 the admixture event occurred 15 generations ago under a somewhat weaker assortative mating (*P* = 0.6), while the estimated number of generations would be 11 under random mating.

**Figure 3:**
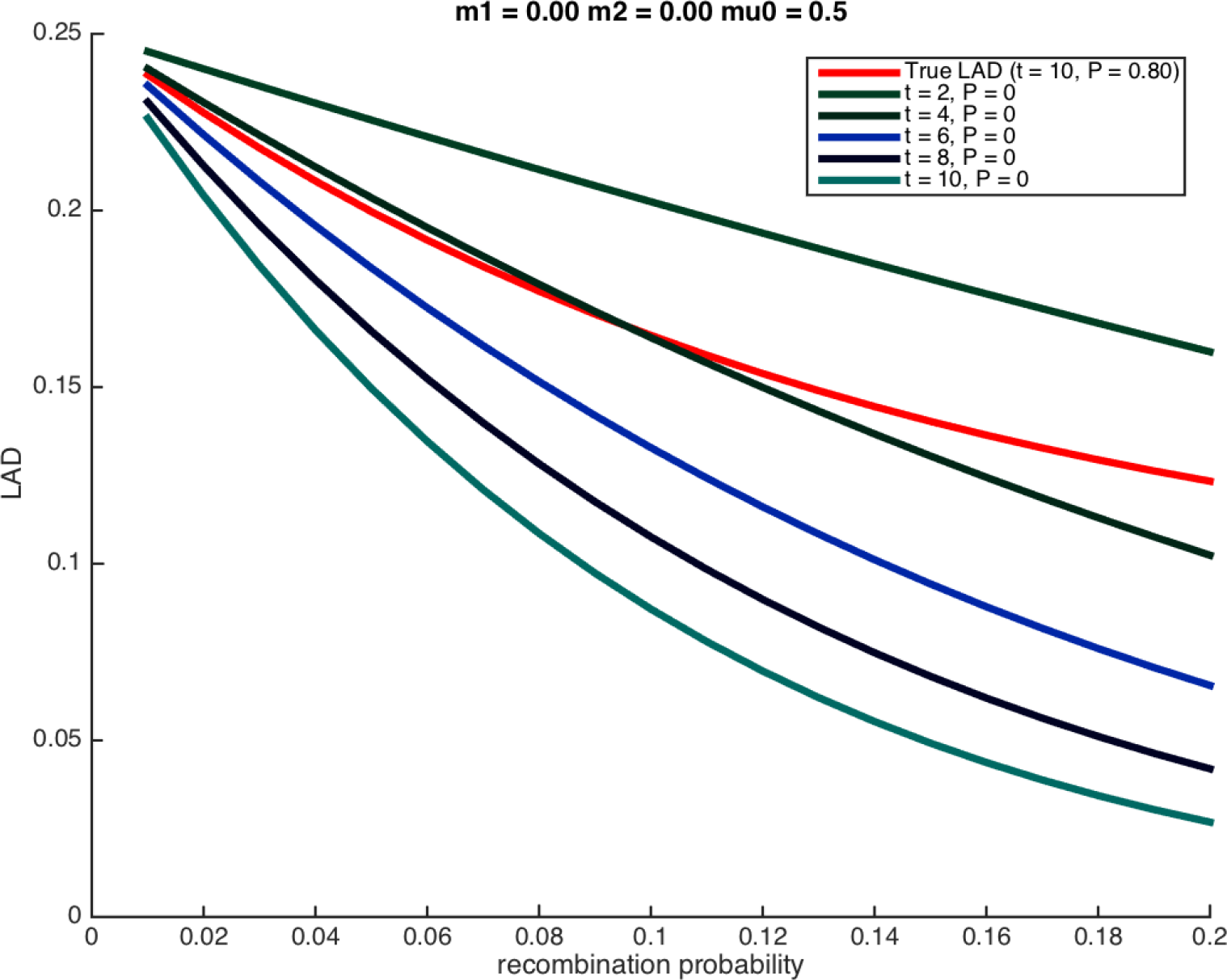
The distribution of LAD for different values of *P* with no migration. The thick lines correspond to the expected LAD based on Lemma 3.1, and the thin lines correspond to simulation runs of a single locus in the genome.

**Figure 4:**
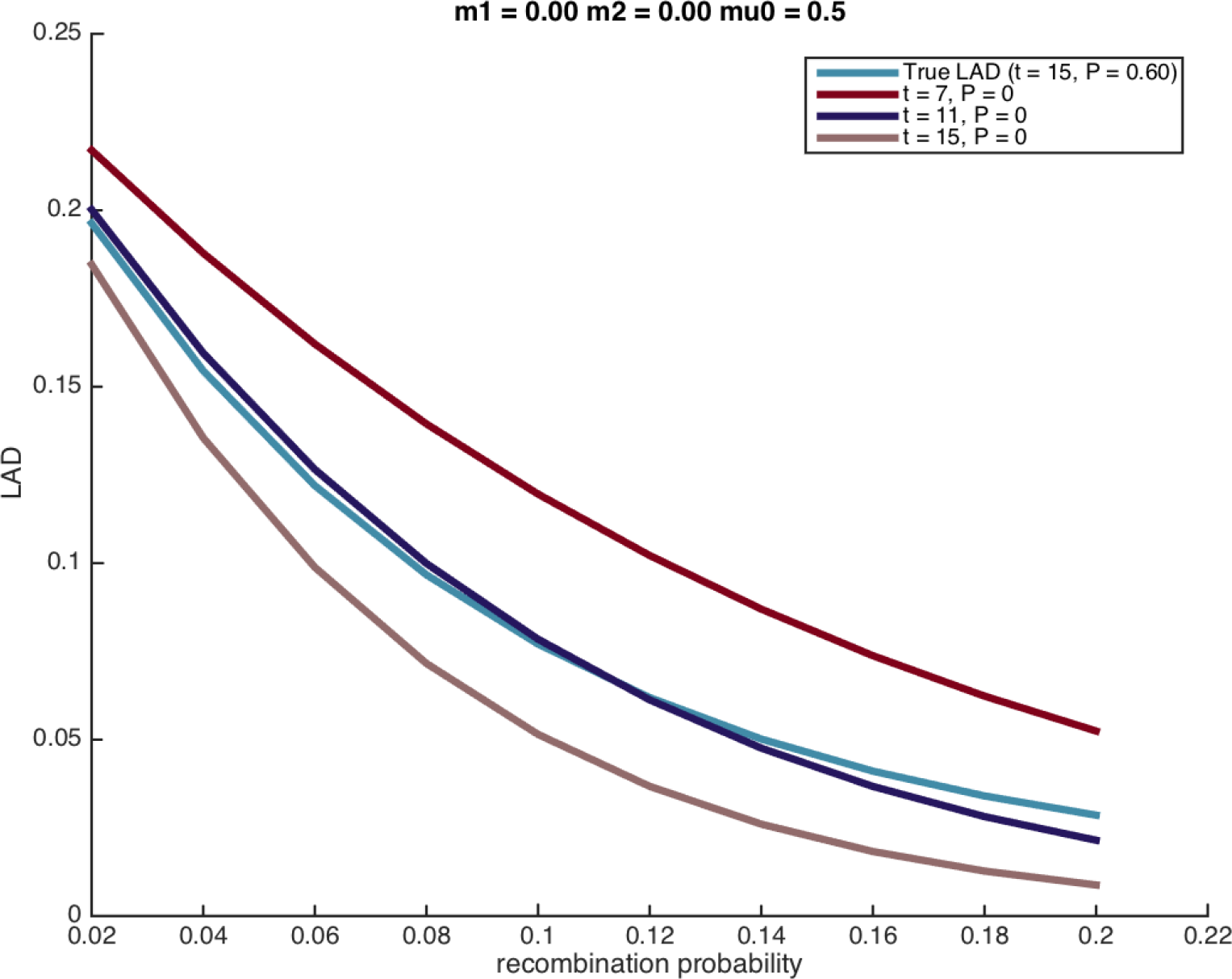
The distribution of LAD for different values of *P* with no migration. The thick lines correspond to the expected LAD based on Lemma 3.1, and the thin lines correspond to simulation runs of a single locus in the genome.

Next, we explore the effect of migration on the LAD function. We consider both the case where the two populations migrate at the same rate (*m*_1_ = *m*_2_) as shown in Figure 5, as well as the case in which *m*_1_ = 0, as shown in Figure 6. Evidently, the theoretical calculations capture the empirical well in the sense that they allow for a clear distinction between different migration rates.

**Figure 5:**
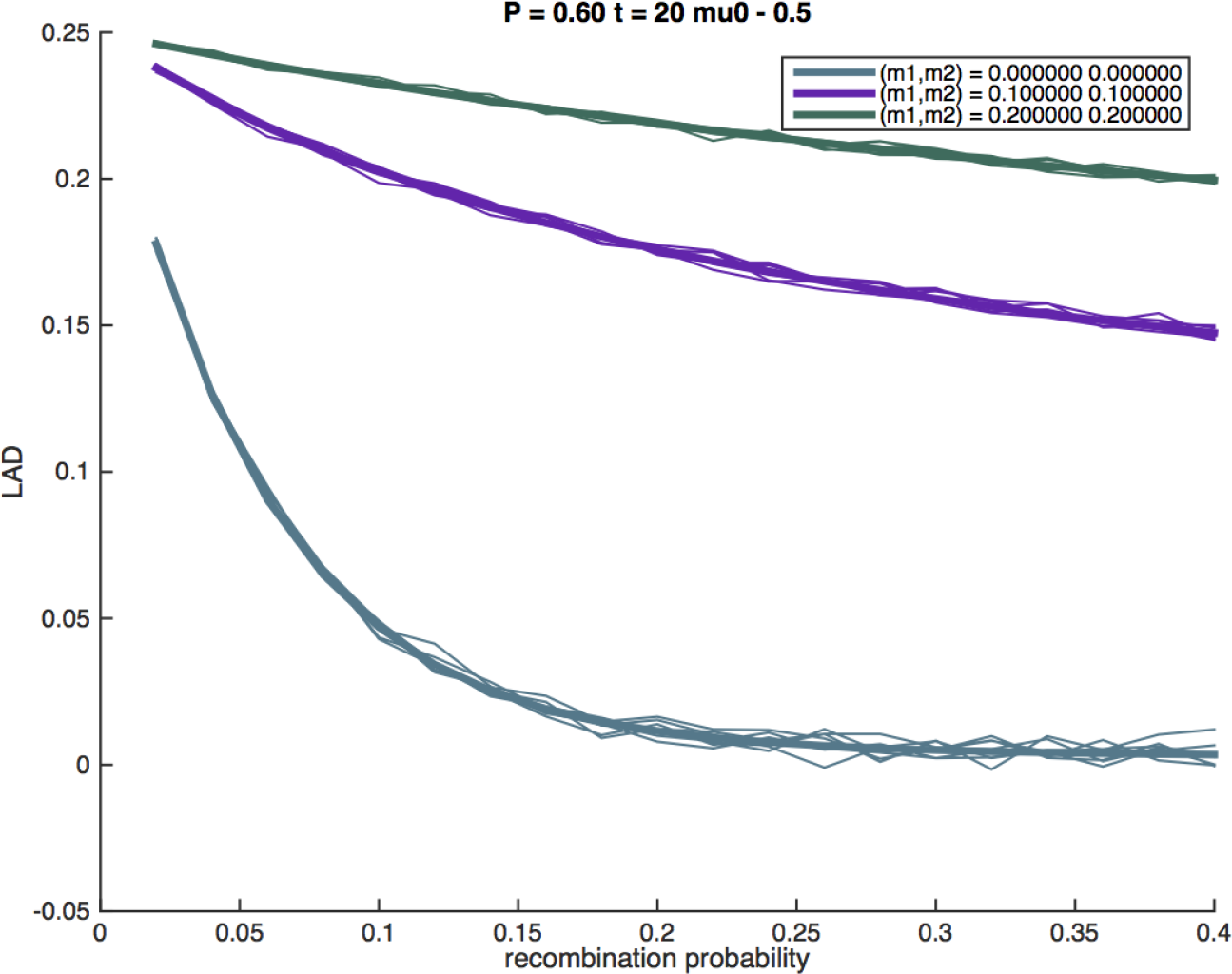
The distribution of LAD for different values of *m*_1_, *m*_2_, with equal migration rates from both populations. The thick lines correspond to the expected LAD based on Equation 2, and the thin lines correspond to simulation runs of a single locus in the genome.

**Figure 6:**
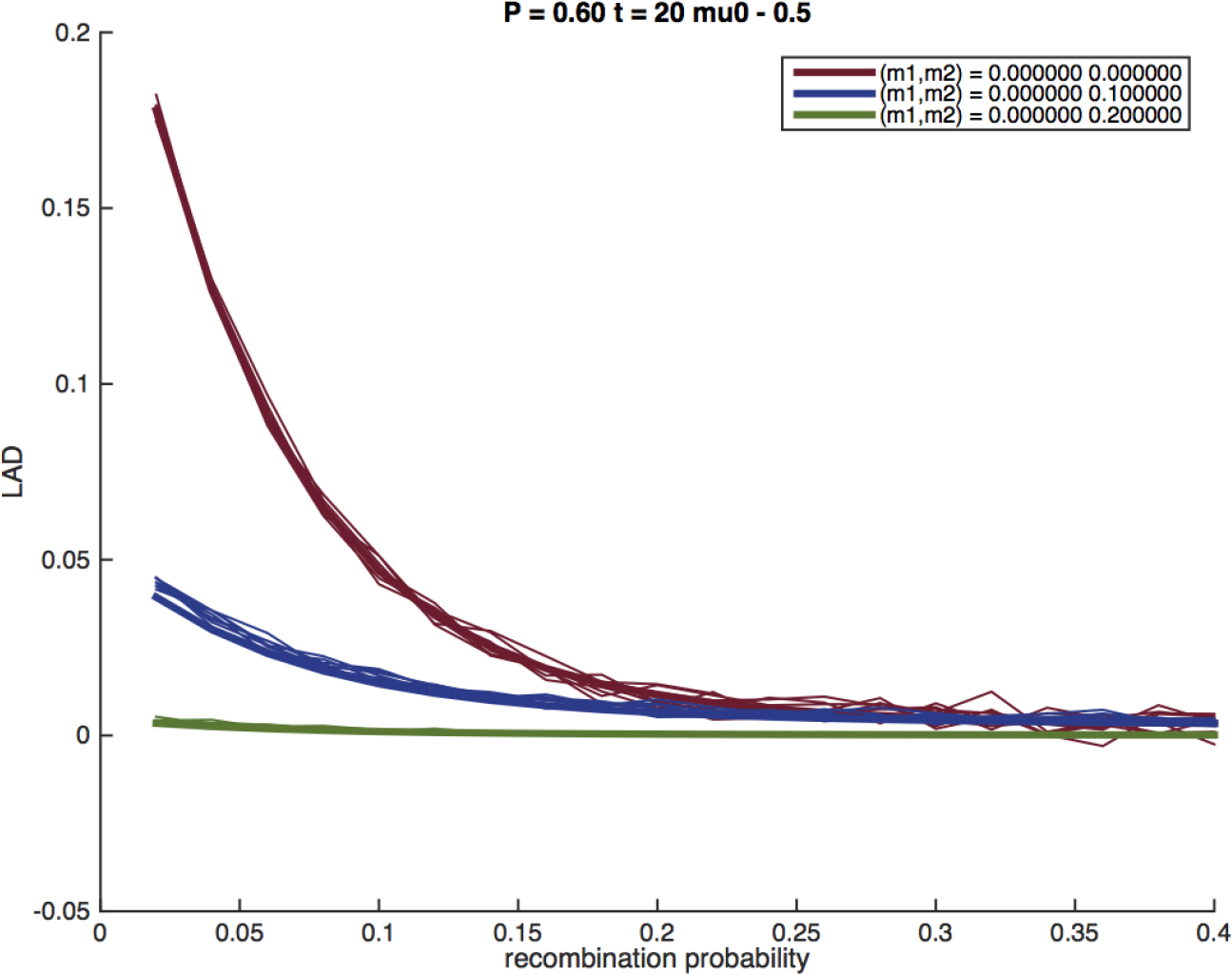
The distribution of LAD for different values of *m*_1_, *m*_2_, with no migration from population 1. The thick lines correspond to the expected LAD based on Equation 2, and the thin lines correspond to simulation runs of a single locus in the genome.

We note that migration and assortative mating can result in similar LAD decay. We estimated the LAD curve using the formula of Lemma 3.1 under random mating with migration, as well as under assortative mating with different values of migration. Since the parameter space (*m*_1_,*m*_2_, *P*) is large, there are triplets of values with very similar LAD curves, thus in practice the model parameters will not necessarily be identifiable. In Figure 7 we present an example where identifiability requires the comparison of LAD decay over dozens of megabases.

**Figure 7:**
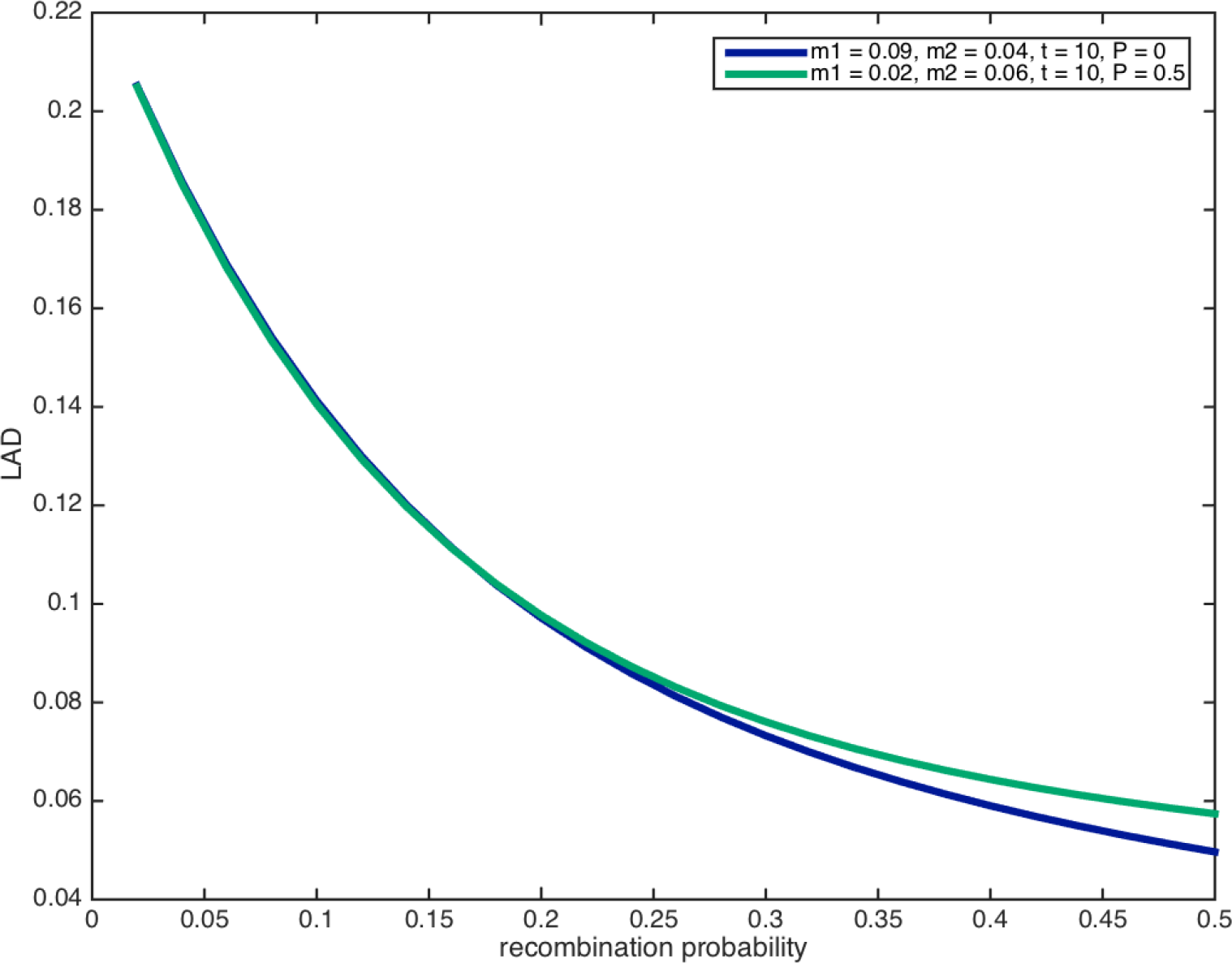
The expected LAD decay under two conditions, one with assortative mating and another with random mating. In the presence of migration the two curves almost overlap, and distinguishing between the two cases will be challenging in practice, particularly if LAD is measured only up to a few dozen centimorgans.

**Results on real data** To examine the properties of our model in real data we used genetic data from 1730 African-American individuals from the the Study of African Americans, Asthma, Genes and Environments (SAGE). The individuals in the SAGE data were genotyped at 800,000 SNPs on the Affymetrix Axiom Genome-Wide LAT 1 Array, and genotype calling and QC were performed as previously described [15].

To compute LAD we first called local ancestry using the LAMP-LD software package[16] and genome-wide ancestry was inferred from mean value of local ancestry for each individual. We measured the LAD decay in 164 **10Mb** overlapping windows with a **1Mb** overlap. We calculated the mean LAD decay across all windows as well as the squared distance of each window to the mean. Regions that are under selection or in which the estimates of recombination rates are inaccurate will result in a different LAD decay. We therefore performed additional QC by removing windows with a LAD decay greater than two standard deviations from the mean. We repeated this process until convergence leaving 96 windows.

We measured the assortative mating over the last generation by applying the method ANCESTOR [12] to the data. ANCESTOR takes as input local and global ancestry and determines the ancestral proportions of the mother and the father of each individual. The Pearson correlation coefficient between the parental ancestries was *P* = 0.32 estimated across all individuals. This establishes that there was strong ancestry based assortment in African Americans in the last generation. If this ancestry-based assortative mating exists in previous generations our theory shows that LAD decay will be affected. Under the assumption that this correlation was stable throughout history, one can use this estimate to constrain the potential demographic histories of African Americans inferred via LAD.

We fitted the migration and assortative mating parameters using a grid search over the entire range of parameters. The best fit resulted in an estimate of *t* = 13 generations, with migration rates *m*_1_ = 0.01, *m*_2_ = 0.05, and assortative mating *P* = 0.46 (Figure 8a). Next, we made the assumption of no migration by searching the grid but with the constraint *m*_1_ = *m*_2_ = 0, but we allowed for assortative mating. In this case, the number of generations was dramatically shortened to 8 generations, and the assortative mating value increased dramatically to *P* = 0.6 (Figure 8b). Similarly, we search the grid with the constraint *P* = 0 in order to study the case of random mating with migration. In this case the number of generations was 16, and the migration values slightly increased to *m*_1_ = 0.02, *m*_2_ = 0.05 (Figure 8c). Finally, under random mating and no migration the estimated number of generation is *t* = 3, which is clearly a vast underestimate of the true number based on the known history of African Americans (Figure 8d). Notably, there is no good fit under random mating and no-migration, and the best fit is obtained in the presence of both migration and assortative mating.

**Figure 8:**
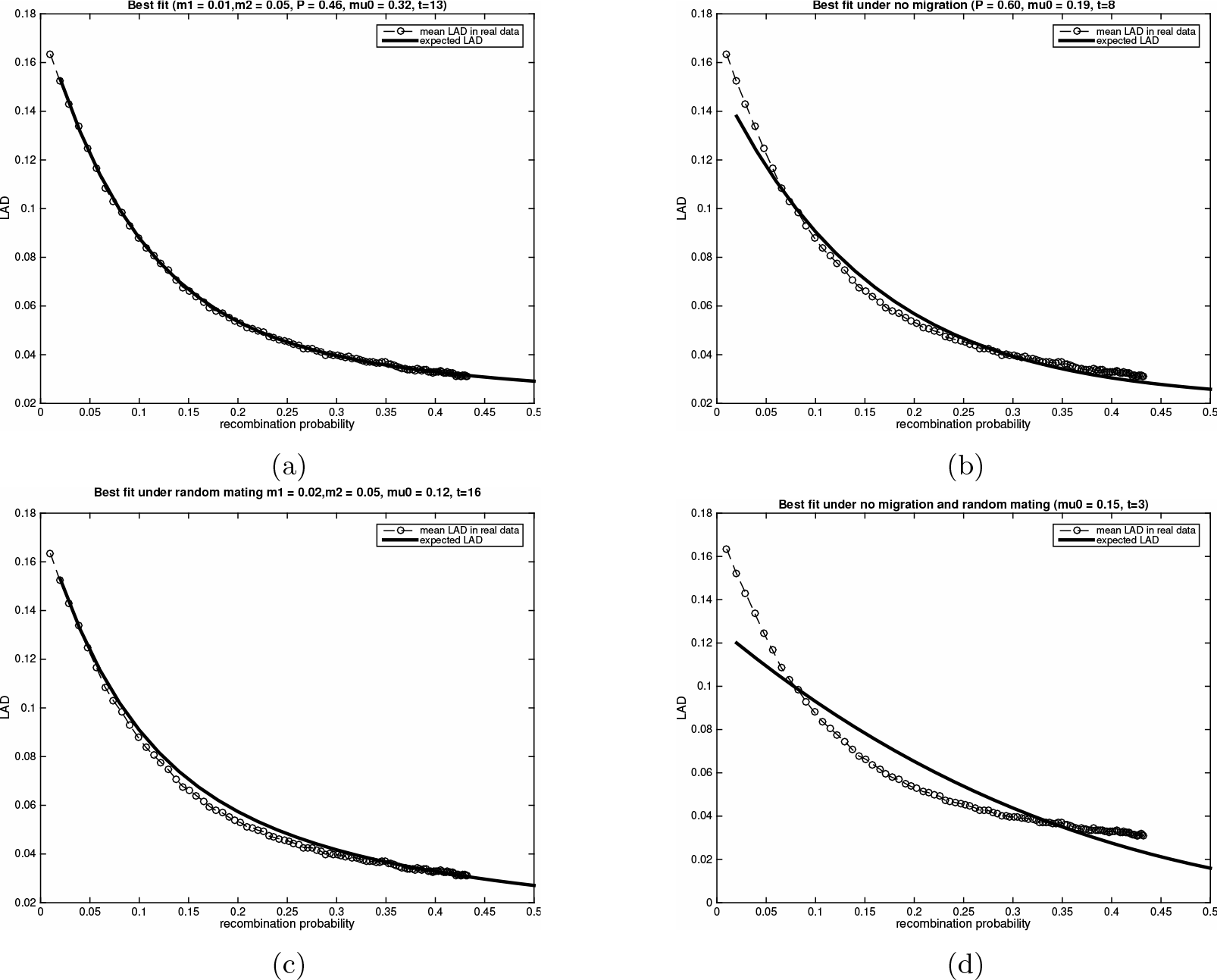
Each of the plots shows the best fit of the parameters to the mean LAD in the African American SAGE dataset: (a) The parameters searched over the entire grid, resulting in the best fit with estimated number of generations 13, migration rates *m*_1_ = 0.01, *m*_2_ = 0.05, and correlation *P* = 0.46. (b) The best fit under the assumption of no migration. The number of generations estimated to be 8, and *P* = 0.6. (c) The best fit under the assumption of random mating with migration. The number of generations is estimated as 16. (d) The best fit under the assumption of random mating and no migration - the number of generations is estimated as 3.

Clearly, the LAD decay is only one summary statistic that depends on the parameters *m*_1_, *m*_2_, *t*, *P*, and other statistics may give somewhat different results. For example, it may be possible to examine the distribution of IBD [9], local ancestry [8], and LD [7] under an assortative mating model. Moreover, the LAD decay is not identifiable since different sets of parameters often lead to similar LAD decay. Particularly, in the case of the African Americans in SAGE, the best fit was followed by a few different sets of parameters. Particularly, under the assumption that *P* = 0.32 is fixed across the generations, the best fit was with *t* = 15 generations, and the migration rates were *m*_1_ = 0.08, *m*_2_ = 0.01. Due the computational complexity of the grid search used to estimate model parameters it was not feasible to estimate confidence intervals. However, as was the case in simulations, migration rates and generation times could be altered to accommodate removing assortative mating form the model.

All genetic data are available via dbGAP with the accession number phs000355.v1.p1.

## 5 Discussion

We presented an adaption of the Wright-Fisher model, which incorporates ancestry-assortative mating in admixed populations. We demonstrated that under this model the linkage disequilibrium of local ancestry (LAD) between markers is a function of their recombination rate, the ancestral population migration rates, and the strength of ancestry based assortment. Assortative mating is likely impacting other estimates of population and medical genetic parameters both within admixed and continental populations including identity by descent distributions, estimates of heritability, joint site frequency spectra, runs of homozygosity, and the distribution of local ancestry track lengths.

While the focus of this work is the definition and presentation the ancestry-assortative model and its properties, we also estimated the parameters of the model in a real African-American data set. Our estimate of 15 generations since admixture in African Americans is larger than previous estimates[8, 9] and it fits considerably better the known history of African Americans[13]. This suggests that taking assortative mating into account may in some cases be critical in order to obtain the correct demographic history or other population parameters.

The approach we presented for estimating the number of generations since admixture using LAD has its limitations. First, this approach involves a very inefficient grid search, resulting in an inability to provide errors around estimates via bootstrap. Second, in some cases both migration and assortative mating can give rise to similar LAD distributions, and therefore in those cases one can mistakenly believe that the migration is higher and assortative mating is lower or vice versa. The latter, however, raises an interesting question: In previous attempts for learning demographic histories of humans and other species, is it the case that the migration coefficients were inflated, or number of generations since admixture deflated due to assortative mating?

Going forward it will be interesting to determine if assortative mating has biased other recent estimates of demographic events such as the introgression of Neanderthals [17] or the domestication of dogs and pigs[18, 19]. We will also explore extensions to multi-way admixed populations and the use of MCMC to provide confidence intervals for parameter estimates. In addition to altering the distribution of LAD, we showed that assortative mating increases the variance of global ancestry. Under certain polygenic models this will induce a concomitant increase in phenotypic variance, which may have implications for selection and evolution.

Our method makes several strong assumptions, which are likely incorrect, such as constant ancestry-assortment strength and migration rates. However, these are a relaxation of previous methods, since for example under the standard Wright-Fisher model, both random mating and no migration are assumed, and thus both migration rates and ancestry-assortative strengths are fixed across the generations in this case (fixed with value 0). While assortative mating has been well studied, to the best our knowledge this is the first attempt to include ancestry-assortment in the estimation of demographic histories. We also reported, for the first time, the strength of ancestry-assortment in African Americans in the previous generation. In future work we intend to examine the effect of ancestry assortment on other genetic features as well as the resulting impact in population and medical genetics.

## 6 Acknowledgments

The authors acknowledge the patients, families, recruiters, health care providers and community clinics for their participation. In particular, the authors thank Sandra Salazar for her support as the SAGE II study coordinator. This work was supported in part by the Sandler Foundation, the American Asthma Foundation, the RWJF Amos Medical Faculty Development Program, Harry Wm. and Diana V. Hind Distinguished Professor in Pharmaceutical Sciences II, and the National Institutes of Health (ES015794 and MD006902). NAZ was supported by an NIH career development award from the NHLBI (K25HL121295). EH was supported by the Israel Science Foundation (Grant 1425/13), United States-Israel Binational Science Foundation (Grant 2012304), German-Israeli Foundation (Grant 1094-33.2/2010) and by the National Science Foundation (Grant III-1217615). The SAGE study was supported by the Sandler Family, American Asthma Foundation, NIH/NIHMD- 1P60 MD006902, NIH/NHLBI - 1R01HL117004-01, NIH/NIEHS - R21ES24844-01, NIH/NIHMD - U54MD009523, and TRDRP 24RT-0025.

